# Dual-light photodynamic therapy administered daily provides a sustained antibacterial effect on biofilm and prevents *Streptococcus mutans* adaptation

**DOI:** 10.1101/2020.01.09.899963

**Authors:** S Nikinmaa, H Alapulli, P Auvinen, M Vaara, J Rantala, E Kankuri, T Sorsa, J Meurman, T Pätilä

**Author notes:** Address for correspondence: Tommi Pätilä, MD, PhD, Otakaari 5 I 436, 02150 Espoo, Aalto University, Finland.

## Abstract

**Introduction:** Antibacterial photodynamic therapy (aPDT) and antibacterial blue light (aBL) are emerging treatment methods auxiliary to mechanical debridement for periodontitis. APDT provided with a near infrared (NIR) light in conjunction with an indocyanine green (ICG) photosensitizer has shown efficacy in several dental in-office-treatment protocols. In this study, we tested *Streptococcus mutans* biofilm sensitivity to either single-light (aPDT or aBL) or dual-light aPDT (simultaneous aPDT and aBL) exposure.

**Materials and Methods:** Biofilm was cultured by pipeting diluted *Streptococcus mutans* suspension with growth medium on the bottom of well plates. Either a single-light aPDT (810-nm aPDT or 405-nm aBL) or a dual-light aPDT (simultaneous 810-nm aPDT and 405nm aBL) was applied, in both cases together with the ICG photosensitizer, while keeping the total light energy constant at 100J/cm^2^. Single-dose light exposures were given after one-day, four-day or fourteen-day biofilm incubations. Also, a repeated daily dose of the same light energy was applied during biofilm incubations on four-day and fourteen-day biofilms. Finally, antibacterial action of the dual-light aPDT with different relative ratios of 810 nm and 405 nm of light energy was examined on the single-day and four-day protocols. Biofilms were scraped, diluted into ratios between 1:1 to 1:100 000, and plated. After re-incubation, colony-forming units (CFUs) were counted, and confocal 3D biofilm imaging was performed.

**Results:** On a one-day biofilm, dual-light aPDT was significantly more efficient than single-light aBL or aPDT, although all modalities were bactericidal. On a four-day maturated biofilm, a single exposure of aPDT or dual-light aPDT was more efficient than aBL, resulting in a four logarithmic scale reduction in bacterial counts. Surprisingly, when the same amount of aPDT was repeated with a daily dosing on a four-day or a fourteen-day biofilm, bacterial viability improved significantly. A similar but milder response of improved bacterial viability was seen after repetitive aBL application. The viability improvement was eliminated when dual-light aPDT was applied. By changing the relative light energy ratios in dual-light aPDT, a relative increase in aBL improved the antibacterial action of dual-light aPDT when the biofilm was older.

**Conclusion:** When aPDT is administered repeatedly to *S. mutans* biofilm, a single wavelength-based aBL or aPDT leads to a significant biofilm adaptation and increased *S. mutans* viability. The combined use of aBL light in synchrony with aPDT arrests the adaptation and provides significantly improved and sustained antibacterial efficacy.

## Introduction

Oral hygiene based on mechanical cleansing by removal of the biofilm has been proven to be the best method for the prevention of dental and periodontal disease [1]. While saliva contains some 700 different bacterial species, regularly performed mechanical biofilm removal essentially only leaves early-forming Streptococcal residual biofilm on the dental surface [2]. Dental and periodontal diseases result from prolonged biofilm infections[3]. Neglected hygiene is usually a factor in the complex multispecies biofilm required for the disease process. While caries essentially involves mainly gram-positive, *Streptococcus*-rich and carbohydrate fermenting biofilm, gingivitis and periodontitis are related to gram-negative, proteolytic bacterial biofilm flora[4]. Sometimes genetic or environmental circumstances, such as virulent bacterial strains, may predispose one to disease despite reasonable dental hygiene[5].

Antibacterial photodynamic therapy (aPDT) and antibacterial blue light (aBL) have emerged as solutions for attacking dental biofilm [6,7]. These methods are based on light photon absorption by chromophores, leading to electron transfer reactions that ultimately result in the production of reactive oxygen species (ROS). Similarly, ROS are used in bacterial killing by polymorphonuclear leucocytes in the phagocytotic bacterial elimination process [8]. Antibacterial PDT combines an externally provided photo enhancer with a specific light wavelength to excite the nearby oxygen into a singlet state, which is mostly responsible for the antibacterial effect. Antibacterial blue light, however, is based on the same mechanism, but the electron transfer reaction occurs by inherent photosensitizers found within the bacteria themselves, mostly porphyrins and flavins. Certain Streptococcus species, including *S. mutans*, are vulnerable to aPDT due to poor ROS scavenging capacity, mostly reflecting the lack of catalase enzyme. On the other hand, aBL is particularly effective for those bacteria carrying the most abundant amount of blue-light absorbing porphyrins, such as the cell-surface black pigment, iron protoporphyrin IX, in the so-called black-pigmented bacteria group [9,10].

Indocyanine green (ICG) is a widely used aPDT photosensitizer in dentistry, due to its low toxicity, non-ionizing properties, water solubility and light absorption at near-infrared (NIR) wavelengths, which have a good tissue penetration. Several studies have shown the efficacy of NIR 810 nm/ICG aPDT as an adjunctive periodontal treatment. In these studies, dosing has been infrequent, mostly due to the light administration requiring a dentist’s in-office equipment and expertise. Auspiciously, a rapid development in light emitting diode (LED) technology has allowed for the development of personal products for light application used at home. In a home setting, the aPDT treatment can be self-administered by the patients themselves, on a more regular and frequent basis.

The antimicrobial efficacy of aPDT is evident for planktonic bacteria, whereas in biofilms, bacteria are more resistant to any antibacterial treatment [11–13]. Furthermore, data on the effect of repetitive aPDT is sparse. Generally, bacteria are unable to produce resistance against aPDT and aBL, although some adaptation has been shown [14–16]. In this study, we tested the efficacy and effect sustainability of aPDT and aBL in an *S. mutans* biofilm-model allowing for repeated daily treatment administrations. We compared single-light aPDT or aBL head-to-head and assessed the effect of their simultaneous application, called dual-light aPDT treatment. We also examined the bactericidal effect of different ratios of aPDT and aBL when the dual-light aPDT was applied. Finally, to analyse the aPDT action mechanism, we evaluated ICG adhesion to *S. mutans* bacteria, using absorption spectroscopy.

## Materials and methods

Monospecies *S. mutans* biofilm model experiments were performed to study the effect of recurring photodynamic therapy during the biofilm formation process. A minimum of six biofilms for each experiment were grown in surface-treated, flat-bottom Nunclon Delta well plates (Thermo Fisher Scientific Inc, US), which have widely been used for *in vitro* biofilm formation of multiple different bacteria species [10,13,17], specifically, the *S. mutans* species [17–19]. The biofilm experiments were divided into different setups based on biofilm maturation age and the therapy given.

### Study protocols

Applications of both single- and dual-light therapies were scheduled to mimic the daily use of antibacterial light therapy at home for hygiene purposes. In all the experiments, the dosing was kept the same, meaning the total amount of light irradiance, the respective light energy applied and the concentration of the ICG photosensitizer (if given) were identical. In the first setting, *S. mutans* biofilm was incubated for either one day or four days, and a single dose of light with ICG was applied at the end of the growth period. In the second setting, the biofilm was incubated for either four days or fourteen days, and a daily dose of light ICG with was applied during incubation. In each setting, the last treatment was followed by plating onto brain heart infusion (BHI)-agar dishes for colony-forming unit (CFU) counting.

### Biofilm model

*Streptococcus mutans* (ATCC 25175) bacteria were grown for 18 h in an incubator (NuAire DH autoflow 5500, NuAire inc, US), at +36 degrees C, 5% CO_2_ in BHI broth (Bio-Rad 3564014, Bio-Rad Laboratories, Inc, US). The resulting bacterial suspension was diluted with a 0.9% NaCl solution until an optical density (OD) of 0.46 was reached. The optical density was measured by a spectrophotometer (Varian Cary 100 Bio UV-VIS, Agilent Technologies, Inc, US), and then with a Den 1 McFarland Densitometer (Biosan, Riga, Latvia).

Biofilms were grown in flat-bottom 96-well plates (Thermo Fisher Scientific Inc, US) by placing 100 μl of 0.46 OD *S. mutans* suspension in each well, with 100 μl of BHI-broth growth medium. The well plates were then incubated in a growth chamber (36°C, 5% CO^2^). The BHI-broth medium was changed daily to supply fresh growth medium and to wash away the debris. The change of the medium in each well was performed by removing 100 μl of the medium and replacing it with a similar amount of fresh BHI broth.

### Light Exposure

Before the light exposure took place, the growth medium was meticulously removed by pipetting and subsequently replaced with an equal amount of indocyanine green solution (Verdye, Diagnostic Green, GmBH), tittered to a concentration of 250 μg/ml. The indocyanine green was left to incubate at room temperature and in the dark for 10 minutes. After this incubation period, the biofilm was washed with a 0.9% NaCl solution. Then, the 0.9% solution of NaCl was added to each well to reach a total volume of 200 μl. Light exposure was performed using specific, custom-made LED light sources (Lumichip Oy, Espoo, Finland). The exposure time was calculated from the determined light amount and known irradiances, which had been previously measured with a light energy meter (Thorlabs PM 100D with S121C sensor head, Thorlabs Inc, US) and a spectroradiometer (BTS256, Gigahertz-Optik GmbH, Germany), respectively. After the exposure, the BHI broth was changed and the plates were placed in the incubator, or, if the light exposure was final, the biofilm was scraped for CFU counting, as described below. Excitation lights were applied with two single-wave LED light sources with peak intensities at 810 nm or at 405 nm, and with a dual-wave LED light chip simultaneously producing two separate peak intensities at 405 nm and at 810 nm.

Antibacterial photodynamic therapy light exposure was administered at an 810-nm peak wavelength LED array on top of the well plate. The resulting light irradiance was 100mW/cm^2^, and the provided light energy was 100J/cm^2^. Antibacterial blue light was administered at a 405-nm peak wavelength LED array, with a resulting irradiance of 80mW/cm2, and resulting light energy of 100J/cm^2^. The dual light was administered with two light peaks identically placed and providing LED arrays on top of the well plate, producing a synchronous irradiance of 50mW/cm^2^ for the 405-nm light, and 50mW/cm^2^ for the 810-nm light. The light energies produced were 50J/cm^2^ (405 nm) and 50J/cm^2^ (810 nm), respectively. To rule out an LED well-heating effect on bacterial viability, temperature controls were measured (Omega HH41 Digital Thermometer, Omega Engineering, US) in the biofilm wells to confirm temperature levels below 35 degrees during the treatment, with a 100mW/cm^2^ radiant flux.

We also tested the antibacterial efficacy of dual-light treatment in terms of the relative amounts of 810-nm and 405-nm light irradiance. Three different light combinations were employed, with a simultaneous use of the single-peak-emitting light sources. Firstly, a 1:1 irradiance ratio of aBL to aPDT was applied, with 70 mW/cm^2^ irradiance for the 405-nm light and 70 mW/cm^2^ for the 810-nm light, the light energy emitted being at 50J/cm^2^ and at 50J/cm^2^, respectively. Secondly, a 3:1 irradiance ratio of aBL to aPDT was applied, with 130 mW/cm^2^ irradiance for the 405nm light and 40mW/cm^2^ for the 810nm light, the light energy provided being at 75J/cm^2^ and at 25J/cm^2^, respectively. Thirdly, a 1:3 irradiance ratio of aBLto aPDT was applied, with 40mW/cm^2^ irradiance for the 405-nm light and 130mW/cm^2^ for the 810nm light, the light energy applied being at 25J/cm^2^ and at 75J/cm^2^, respectively.

### Colony-Forming-Unit Counting

Subsequent to the final light exposure of each experiment, the entire biofilm was removed from the well by collecting the broth together with the dense biofilm. The dense biofilm was mechanically scraped from the bottom of the well plate using a sterile inoculation rod and then placed into a 1-ml test tube, forming 200 μl of suspension. After meticulous vortexing (Vortex Genie, Scientific Industries Inc, US), serial dilutions from 1:1 to 1:100 000 were done, using sterile ART filter tips (Thermo Scientific, Waltham, US). To enumerate the viable cells, 100 μl of resulting biofilm dilution was then evenly spread over an entire BHI plate, using a sterile L-shape rod.

The plates were then assembled into the incubator, the bacteria were grown for 48 h, and the plates were photographed (Canon D5 DSLR camera with Canon EF 24-70 mm f/4L lens, Canon, Japan) on a light table (Artgraph Light Pad Revolution 80, Artograph Inc, US). Colony-forming units were assessed with Image J software (National Institute of Health, US). Typically, numbers between 30 and 800 were considered to be in the range where the data was statistically most reliable, and the number of bacterial colonies was calculated accordingly.

### Confocal scanning laser microscope (CSLM)

The structural organization of the biofilm was examined with confocal fluorescence imaging with a Leica TCS CARS SP 8X microscope (Leica Microsystems, Wetzlar, Germany), using HC PL APO CS2 20X/0.75 numerical-aperture multi immersion and HX PL APO CS2 63X/1.2 numerical-aperture water immersion objectives. The imaged biofilms were stained using a live/dead BacLight bacterial viability kit (Molecular Probes. Invitrogen, Eugene, Oregon, USA). The stains were prepared according to manufacturer directions and were left to incubate at room temperature and in the dark for 15 min prior to examination under the confocal scanning laser microscope (CSLM). Light excitation was performed with a two-laser system, a 488-nm Argon laser and a 561-nm DPSS laser, the emission windows configured to exclude the excitation wavelength of the two lasers and to meet the emission wavelength of the live/dead fluorescence marker. The emission window for the 488-nm laser was set at 500 nm - 530 nm, and for the 561-nm laser, at 620 nm - 640 nm.

### Spectroscopic absorption assessment of ICG within a *Streptococcus mutans* pellet

One ml of *S. mutans* suspension with 0.46 OD, corresponding approximately to 100×10^6^ CFUs, was centrifuged for 5 minutes at 8000 rpm (Heraeus Megafuge 1.0, Thermo Scientific, Waltham, US) to the bottom of a 2-ml Eppendorf tube to form a 10-μl pellet. The supernatant was removed and replaced by a 1-mg/ml ICG solution to establish a total volume of 1 ml. The pellet was then mixed into the solution by vortexing for 60 seconds and left to incubate for 10 minutes, after which it was washed twice by centrifugation of the bacteria into a pellet and by replacing and vortexing the supernatant into a 0.9% NaCl solution. This was followed by re-centrifugation at 8000 rpm for 20 minutes. The *S. mutans*-formed pellet was vortexed into a fresh 0.9% NaCl solution to form 0.46 OD, for an absorption spectroscopy analysis with a Gary 100 Bio UV-visible spectrophotometer (Varian Inc., Palo Alto, CA). For comparison purposes, an ICG 4-μg/ml NaCL 0.9% solution was used. The *Streptococcus mutans* 0.46 OD 0.9% NaCl solution suspension was used as a reference sample for the ICG/S.mutans suspension and 0.9% NaCl for the ICG 4 μg/ml NaCL 0.9% solution.

### Statistical analysis

Colony-forming-unit counts were compared with the non-parametric Mann-Whitney U-test, using GraphPad Prism 8 software (GraphPad Software, San Diego, US).

## Results

The one-day *S. mutans* biofilm exhibited significantly reduced viability when exposed to aBL, from a median of 19×10^6^ CFUs (the range being 2.6×10^6^-34×10^6^ CFUs) to a median of 1.65×10^6^ CFUs (the range being 0.08×10^6^-10×10^6^ CFUs) (p=0.0455, Mann Whitney U-test). Antibacterial photodynamic therapy administered with an 810-nm light together with ICG resulted in a markedly better efficiency than did aBL, with a 3 logarithmic scale reduction in alive bacteria counts compared to the control biofilm, showing a median of 1.3×10^3^ CFUs (the range being 0.3×10^3^-47×10^3^ CFUs) (p=0.0025, Mann-Whitney U-test). However, when aBL and aPDT were combined via the dual-light aPDT, the median of alive bacteria decreased to 0 CFU (the range being 0-0.7×10^3^ CFUs) (p=0.0043, Mann-Whitney test). The bactericidal effect of dual-light aPDT was significantly more efficient when compared to aPDT or aBL provided separately (p=0.0064, p=0.0022, respectively, Mann-Whitney U-test), see Figure 1.

**Figure 1.**
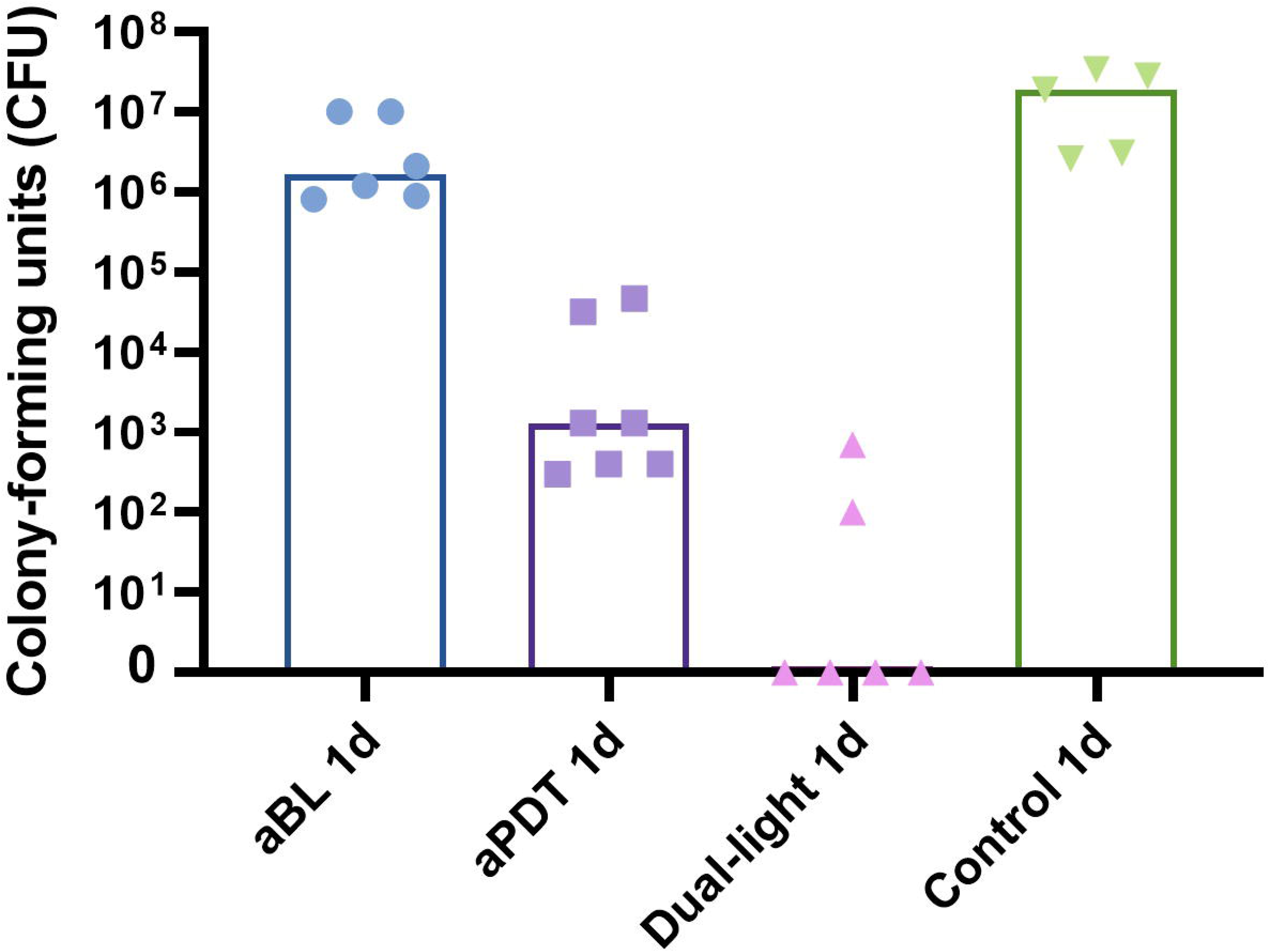
A one-day *S. mutans* biofilm was treated with a single-dose application of aBL, aPDT or dual-light aPDT. The aBL reduced the number of living bacteria significantly, but the antibacterial effect of aPDT was markedly better than that of aBL. The combination of the two, the dual-light aPDT, provided significantly greater antibacterial activity. The total amount of light irradiance was the same at 100mW/cm^2^ for all three modalities (aBL vs. control, p=0.045; aPDT vs. control, p=0.0025; dual-light aPDT vs. control, p=p=0.0043; aBL vs. dual-light aPDT, p=0.0022; aPDT vs. dual-light aPDT, p=0.0064; aBL vs. aPDT, p=0.0012; Mann-Whitney U Test).

In the four-day biofilm model, where a single dose of aBL or aPDT was given at the end of the biofilm maturation period, the aBL reduced the alive bacteria counts to a median of 2.7×10^6^ CFUs (the range being 1.4×10^6^-17×10^6^ CFUs), from a median of 28×10^6^ CFUs (the range being 8.4×10^6^-68×10^6^ CFUs) of the control biofilm (p=0.0245, Mann-Whitney U-test). Again, the aPDT application was significantly more effective than the aBL one, leaving only a median of 2×10^3^ CFUs (the range being 0.1×10^3^-11×10^3^CFUs) (p= 0.0022, aBL vs aPDT, Mann-Whitney test). Similarly to the one-day biofilm test, the simultaneous application of aBL and aPDT in the dual-light aPDT group showed an improved bactericidal effect, leaving a median of 1.9×10^3^ CFUs (the range being 0.1-48×10^3^ CFUs). There was no statistical difference between aPDT and dual-light aPDT (p=0.7381, Mann-Whitney U-test) in the single-dose treatment of the four-day biofilm model. In general, the bacterial viability was better after the single-dose treatment of the four-day biofilm, when compared to the respectively treated one-day biofilms, see Figure 2.

**Figure 2.**
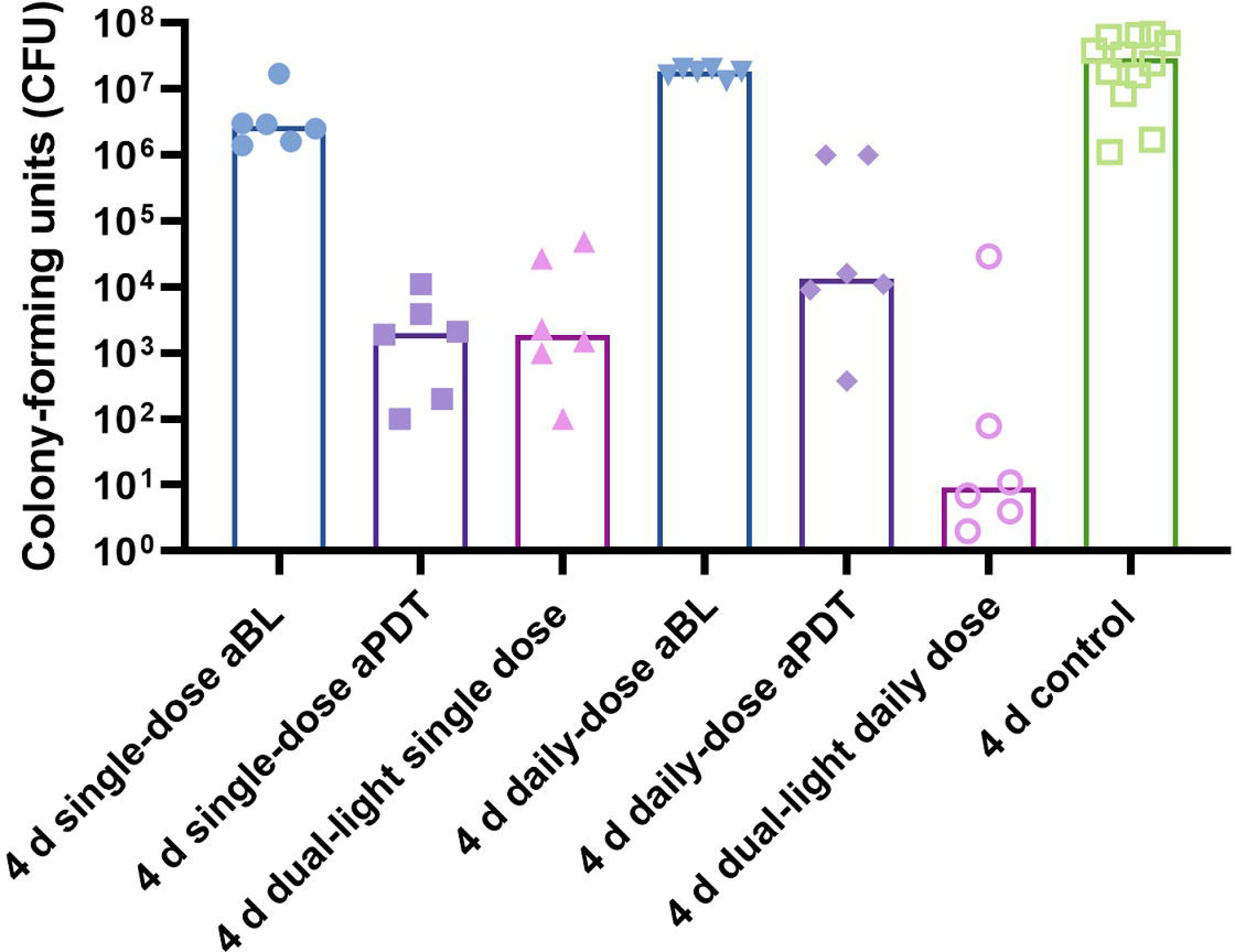
A four-day biofilm was exposed to aBL, aPDT or dual-light aPDT as a single-dose exposure at the end of the biofilm maturation period or as a repetitive, daily-dose exposure repeating the same treatment dose. The columns display medians. Antibacterial blue light decreases the viability of the *S. mutans* biofilm significantly after a single dose at the end of the biofilm maturation period of four days. A daily dose of aBL increased viability of the biofilm, showing no difference to the viability of the control biofilm. The antibacterial effect of aPDT was significantly better than that of aBL, with several ten logarithmic scale increments. However, the viability of the *S. mutans* biofilm improved significantly after four days of the daily-dose aPDT treatment. Single-dose, dual-light aPDT showed a better antibacterial effect when compared to aBL or aPDT alone. More importantly, the antibacterial effect of dual-light aPDT continued to improve after the repeated daily-dose application and reduced the viability of the biofilm significantly (single-dose aBL vs. control, p=0.025; single-dose aPDT vs. control, p=0.0001; dual-light, single-dose aPDT vs. control, p=0.0001; single-dose aBL vs. dual-light, single-dose aPDT, p=0.0022; single-dose aPDT vs. dual-light, single-dose aPDT, p=0.74; single-dose aBL vs. single-dose aPDT, p=0.0022; daily-dose aBL vs. control, p=0.35; daily-dose aPDT vs. control, p=0.0001; dual-light, daily-dose aPDT vs. control, p=0.0001; daily-dose aBL vs. dual-light, daily-dose aPDT, p=0.00022; daily-dose aPDT vs. dual-light, daily-dose aPDT, p=0.026; daily-dose aBL vs. daily-dose aPDT, p=00022; single-dose aBL vs. daily-dose aBL, p=0.0087; single-dose aPDT vs. daily-dose aPDT, p=0.048; dual-light single dose vs. dual-light daily dose, p=0.04; Mann-Whitney U Test)

A daily-dose repetitive application of aBL on the four-day biofilm model showed significantly improved bacterial viability, with a median of 18×10^6^ CFUs (the range being 13×10^6^-20×10^6^ CFUs), when compared to the equivalent dose, i.e. a single-dose application of the aBL to a four-day matured biofilm, as described above (p=0.0087, Mann-Whitney U-test). Similarly, a daily-dose repetitive application of aPDT left a median of 13.5×10^3^ CFUs (the range being 0.38×10^3^-1.0×10^6^ CFUs). This is significantly more than in the four-day biofilm model, where the aPDT was applied as a single-dose treatment (p=0.0476, Mann-Whitney U-test). However, the daily dose of combined aBL and aPDT in the dual-light aPDT group in the four-day biofilm model reduced the alive bacteria to a median of 9 CFUs (the range being 2-29×10^3^ CFUs). Thus, unlike the single-light aBL or aPDT application, the repeated dual-light aPDT application significantly reduced the biofilm viability, when compared to the equivalent dose, i.e. the single-dose application of the same treatment (p=0.0411, Mann-Whitney U-test), see Figure 2.

A daily-dose repetitive application of aBL in the fourteen-day biofilm model showed, similarly to the four-day biofilm model, significantly improved bacterial viability, with a median of 57.5×10^6^ CFUs (the range being 66×10^6^ - 41×10^6^ CFUs), when compared to the four-day biofilm treated with single-dose aBL (p=0.0022), or even the four-day daily-dose repetitive aBL application (p=0.0022). However, the fourteen-day daily-dose of aBL The daily dose of aPDT in the fourteen-day biofilm model similarly improved viability of the biofilm, with a median of 5.1 CFUs (the range being 0.8 - 7.8×10^6^ CFUs), when compared to the four-day biofilm treated by a single dose of aPDT (p=0.0022) or a daily dose of aPDT (0.0087). In the fourteen-day biofilm model, again, the dual-light aPDT outperformed aBL or aPDT, with ongoing improvement in the bactericidal effect, leaving only 1725 CFUs (the range being 1-7.9×10^6^ CFUs), see Figure 3. No significant difference was observed among the dual-light fourteen-day daily-dose biofilm treatment, the four-day single-dose treatment or the four-day daily-dose treatment (p=0.8182 and p=0.4199, respectively).

**Figure 3.**
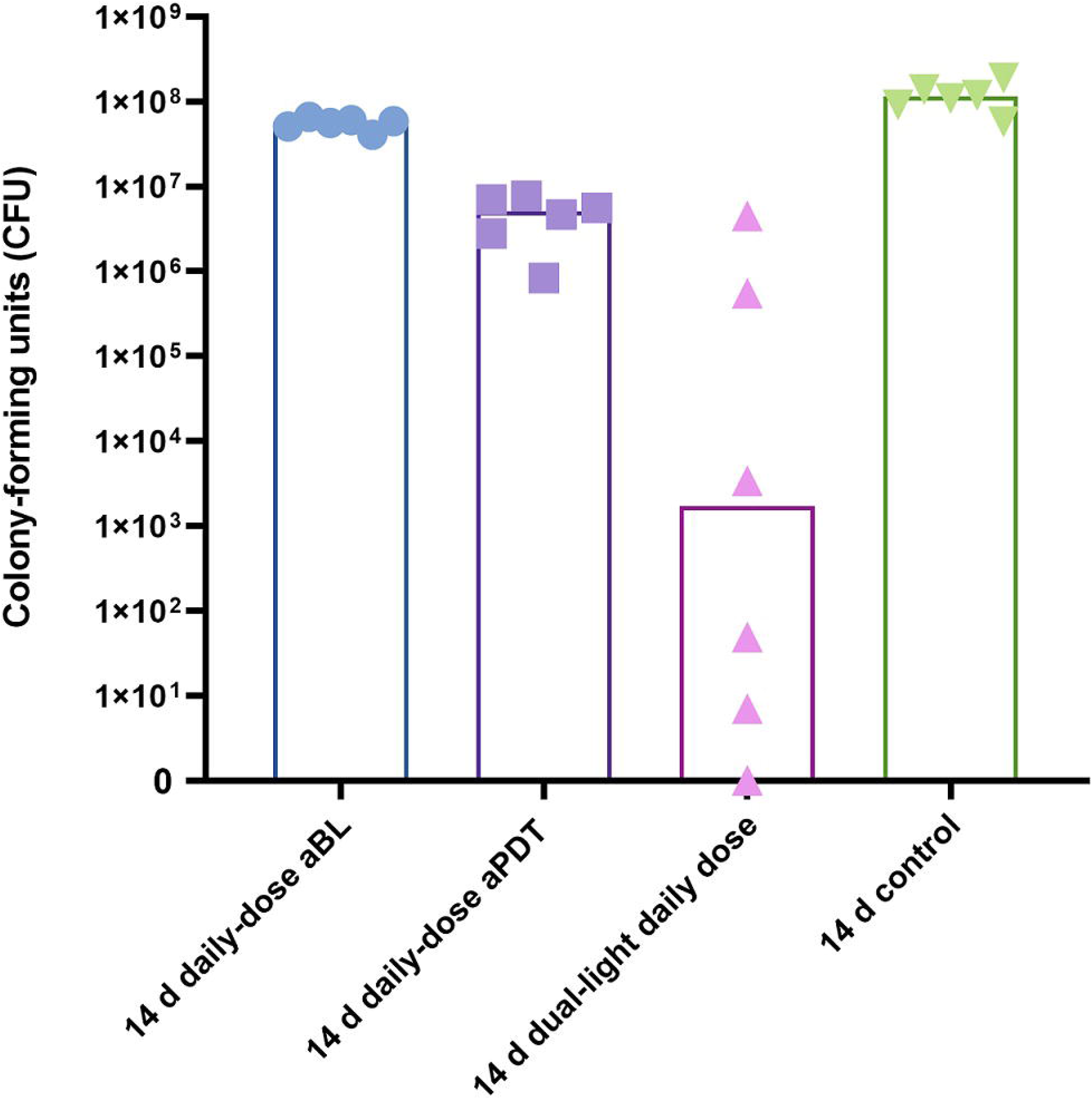
An extended daily-dose study protocol of fourteen days was established to test the ability of the biofilm to adapt to a repetitive aBL and/or aPDT application. A **daily-dose repetitive application of aBL for fourteen days against a *S. mutans* biofilm model showed a reduction in the CFU counts compared to the fourteen-day control biofilm (p=0.02). The CFU counts were significantly more reduced when a daily dose of aPDT was applied (aPDT vs. control, p=0.0022; aPDT vs. aBL, p=0.0022). The dual-light, daily-dose light application showed the most effective antibacterial effect (dual-light vs. aBL, p=0.0022, dual-light vs. aPDT, p=0.0087; dual-light vs. control, p=0.0022, Mann-Whitney U-Test).**

We analysed the impact of relative amounts of aBL and aPDT concerning dual-light aPDT antibacterial efficacy. In the one-day *Streptococcus mutans* biofilm, applying dual-light aPDT at a 1:1 irradiance ratio of aBL to aPDT light provided a median of 0 CFUs (the range being 0-500 CFUs); while a 3:1 irradiance ratio of aBL to aPDT light provided a median of 450 CFU count (the range being 0-7.8×10^3^ CFUs); and the 1:3 irradiance ratio of aBL to aPDT light provided a median of 700 CFUs (the range being 100-25×10^3^ CFUs). In the four-day biofilm model, having a single dose of dual-light aPDT treatment, the 1:1 irradiance ratio of aBL to aPDT light left a median of 100 CFUs (the range being 0-77 10^3^ CFUs); the 3:1 irradiance ratio of aBL to aPDT light provided 10.3×10^3^ CFUs (the range being 0-780×10^3^ CFUs); and the 1:3 irradiance ration of aBL to aPDT provided a median of 0 CFUs (the range being 0-2.2×10^3^ CFUs). Finally, In the daily, repetitive dual-light aPDT application, the 1:1 irradiance ratio of aBL to aPDT light left a median of 0 CFUs (the range being 0-400 CFUs); the 3:1 irradiance ratio of aBL to aPDT light left a median of 2.2×10^3^ CFUs (the range being 0-900×10^3^ CFUs); and the 1:3 light irradiance ratio of aBL to aPDT light left a median of 100 CFUs (the range being 0-740×10^3^ CFUs), see Figure 4.

**Figure 4.**
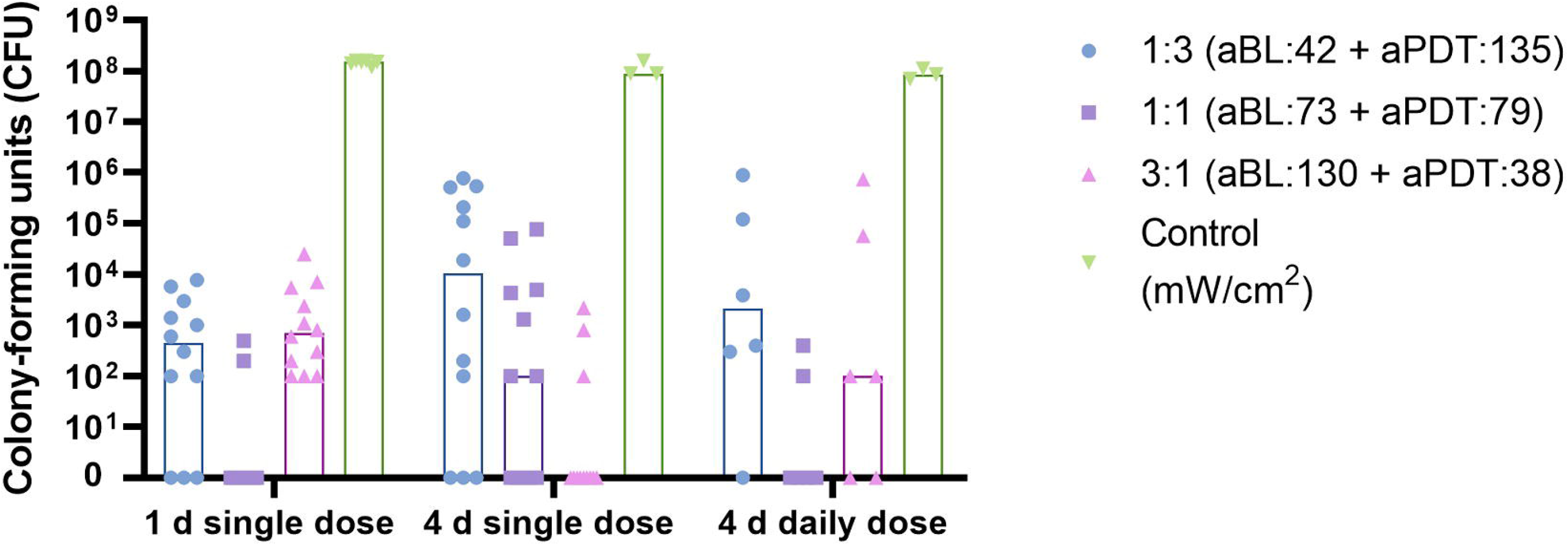
The results of an experiment with dual-light, aPDT-relative-irradiance ratios showed that in the four-day biofilm model, the 1:1 irradiance ratio of aBL to aPDT light had the greatest efficacy of the tested ratios when a daily dose was applied. However, the amount of aBL needed to improve the antibacterial effect of dual light in single-dose treatment is dependent on the age of the biofilm. In an early stage of the biofilm, after a one-day maturation period, the amount of blue light is not so critical, but during maturation of the biofilm, the higher relative amount of blue light seems beneficial, and the 3:1 irradiance ratio of aBL to aPDT ratio showed the most antibacterial effect in the four-day single-treatment protocol. However, the repetitive use of dual light has the most antibacterial effect in the 1:1 setting (single-day, single-dose, dual-light aPDT: 1:3 vs. 1:1, p=0.003; 1:3 vs. 3:1, p=0.43; 1:1 vs. 3:1; p<0.0001; four-day, single-dose, dual-light aPDT: 1:3 vs. 1:1, p=0.12; 1:3 vs. 3:1, p=0.0057; 1:1 vs. 3:1; p=0.067; four-day, daily-dose aPDT: 1:3 vs. 1:1, p=0.045; 1:3 vs. 3:1, p=0.36; 1:1 vs. 3:1; p=0.27; Mann-Whitney U Test).

With CSLM scanning, we were able to visualize the viability of the biofilm after each treatment protocol, see Figure 5. Multidimensional imaging of live (green) and dead (red) bacteria showed most of the live bacteria located at the basal layer of the biofilm, suggesting that this area is more protected against any treatment. In the four-day biofilm model, the surviving bacteria were located at the bottom of a fully developed, thick biofilm. However, when the treatment was applied repetitively for four days, the biofilm appeared visually thinner and denser. The biofilms exposed to repetitive aPDT showed sporadic patchy areas where living bacteria were scattered. These patches could not be found in the dual-light aPDT-treated biofilms, see Figure 6.

**Figure 5.**
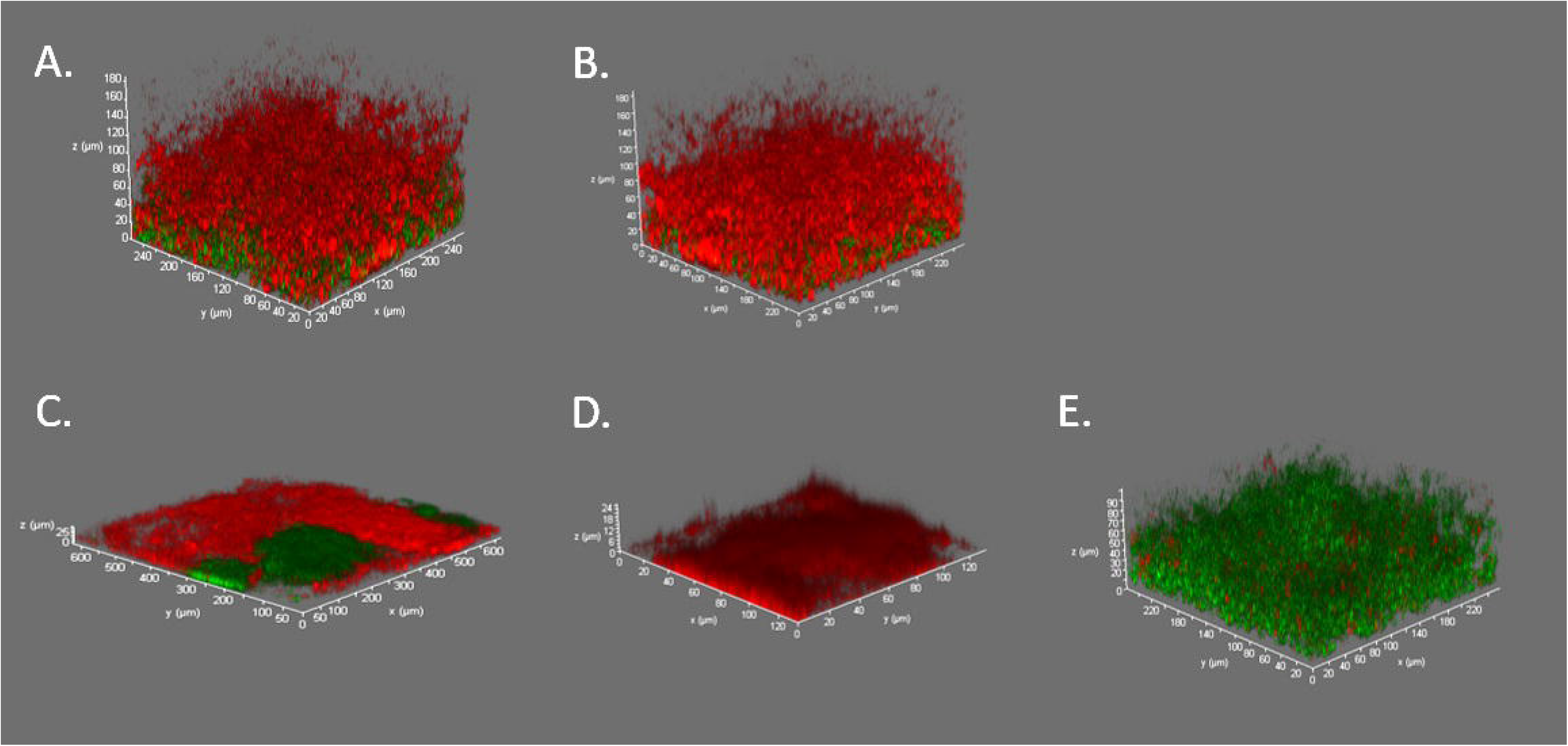
Confocal 3D images of the four-day maturated biofilms stained with live/dead bacterial staining. A. A single-dose application of aPDT significantly reduced the amount of alive *S. mutans* bacteria (green) in the biofilm. The surviving bacteria were mostly located at the basal layer of the biofilm, while most of the dead bacteria (red) were located at the more superficial layer. B. When dual-light aPDT was applied, the viability was significantly less than in the aPDT treated biofilms (also see Figure 2.). C. A daily-dose application of aPDT for four days showed improved viability of *S. mutans* compared to the single-dose treatment. Patchy areas of living cells were found clustered within the biofilm structure, while the overall achitecture of the biofilm was flattened and denser compared to the more fluffy appearance of the freely grown four-day biofilm. D. We could not find these patchy viable areas in the dual-light, aPDT-treated biofilms, where the living bacteria were very scarcely scattered. E. A healthy *S. mutans* biofilm shows great viability with only occasional dead bacteria visible.

**Figure 6.**
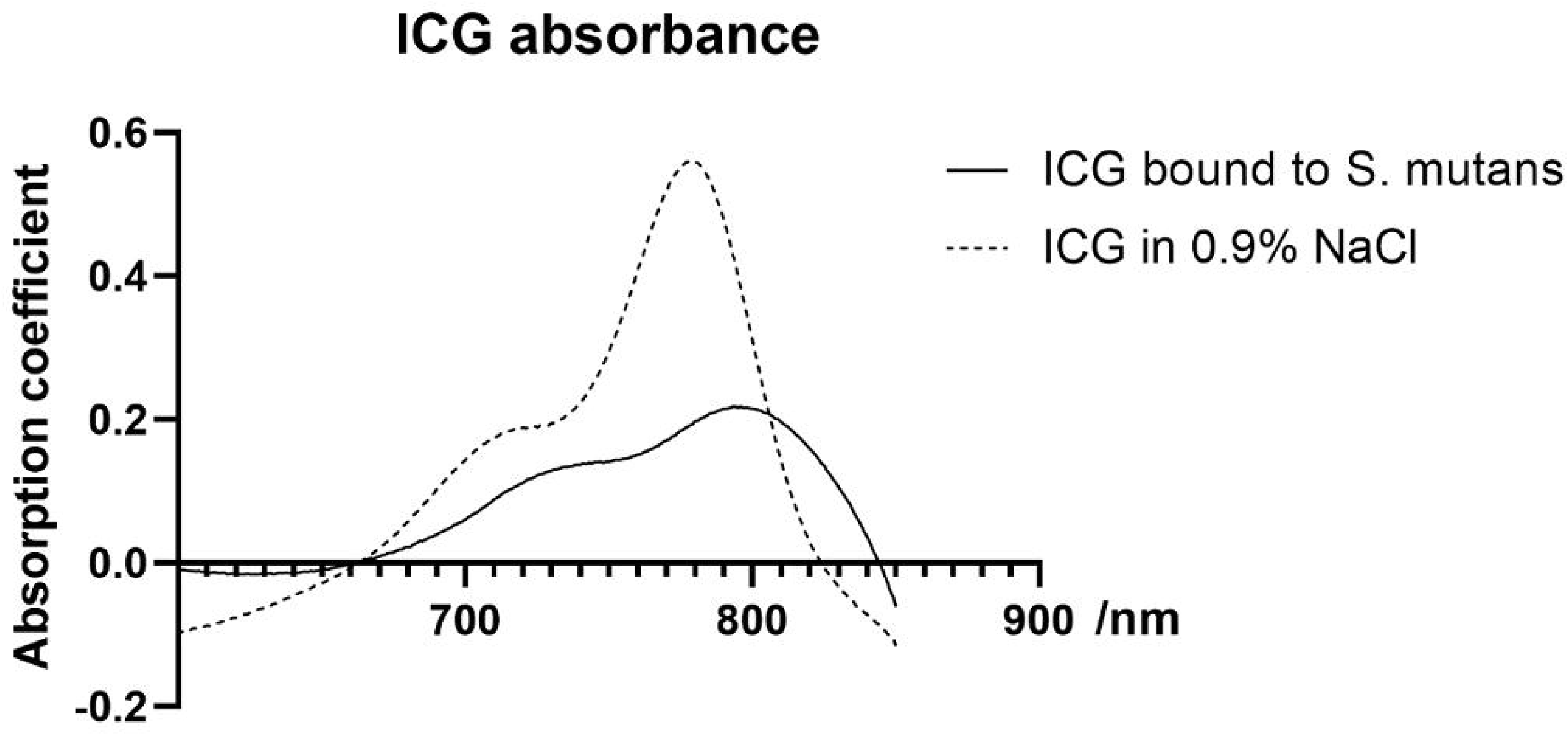
The absorption spectrum of ICG bound to *S. mutans* and the absorption spectrum of free ICG dissolved in 0.9% NaCl.

Bacterial absorption difference between the ICG incubated *S. mutans* cells and the control *S. mutans* suspension in the 0.9% NaCl showed a 20-nm red shift of the absorption spectrum, when compared to the ICG absorption peak in water (see Figure 7). Moreover, the absorption peak was lower.

## Discussion

This is the first study demonstrating the superior efficacy of dual-light aPDT against *S. mutans* biofilm, when compared to single-light aPDT or aBL. The simultaneous, synchronized application of an ICG/810-nm aPDT and 405-nm aBL resulted in a significantly improved antibacterial efficacy, the absolute CFU-count reduction constituting six logarithmic scales, and a persistent antibacterial effect. This persistent antibacterial effect has been nominated as substantivity, when applied to oral hygiene. Such substantivity was not seen in the four-day repetitious aBL exposure, where bacterial CFU counts increased up to about five-fold, when compared to a single-dose aBL application. The ability of *S. mutans* to adapt to the repetitive aBL treatment eventually resulted in a viability comparable to the control biofilm. Similarly, the retained antibacterial action abated when repeated aPDT was applied. Although aPDT showed a significantly better antibacterial effect compared to aBL in *S. mutans* biofilm, the biofilm adapted to repeated exposure with increased CFU counts of up to 100-fold when compared to the single-dose aPDT treatment. The response to the repeated adverse environmental stimuli developed in the very early stage, within the first few days or doses of repeated exposure. The dual-light aPDT thus markedly outperformed both aPDT and aBL in efficacy, but most importantly, the synchronized use was able to suppress the ability of the biofilm to adapt to the external stress. This suppression provided the persistent action required if the method were to be adapted for clinical use in dentistry.

We assessed the relative amount of blue light needed to improve the efficacy of aPDT. The relative increase of aBL in the total light energy of dual-light aPDT decreased the absolute amount of the aPDT effect, because the total amount of light was kept constant. The relative increase in the aBL was effective against older biofilm, showing the 3:1 aBL ratio as being the most effective against four-day old biofilm, but the repetitive dual-light dosing was most effective when the aBL ratio stood at 1:1 with the 810-nm light. Of the different blue light spectrums, we chose to use 405-nm aBL for two main reasons. Firstly, the antibacterial efficacy of 405-nm light has been shown to outperform longer aBL wavelengths in several studies [20]. Secondly, even in the visible light spectrum, there are variations in harmfulness to eyes with different light wavelengths, and eye safety improves at 405-nm light, as compared to 450-nm light or other aBL alternatives.

Different photosensitizers and excitating light combinations have been used against dental biofilms [6]. Indocyanine green, widely used and tested in dentistry, has been approved by the Food and Drug Administration in the United States of America for this purpose. It has been shown as a rather weak singlet oxygen provider, but it does possess a temperature-raising antibacterial ability within a biofilm. After light absorption, an ICG molecule can reach the ground state by releasing the energy through three different pathways. Firstly, the energy can convert into a fluorescence emission ranging from 750 nm to 950 nm. The spectrum maximums are approximately 780 nm in water and 810 nm in blood. Secondly, part of the energy is transferred to an ICG triplet state via intersystem crossing, being able to produce reactive oxygen species. The yield of triplet formation of ICG is 14% in water, and 11% in an aqueous albumin solution. The quantum yield of triplet formation of ICG is sufficiently high for generating efficient reactive oxygen species, particularly singlet oxygen. Thirdly, the energy can be transformed into heat within the ICG molecule itself by internal conversion. It has been estimated that as much as 85% of the absorbed energy could be converted into heat [21]. The ability of ICG to produce antibacterial action through different mechanisms provides an attractive safety feature, especially if aPDT were to be administered frequently. The thermal antibacterial mechanisms could also provide an additional benefit when antibacterial efficacy is provided in deep periodontal pockets, where oxygen in not readily available.

We ruled out the macroscopic LED well heating effect by confirming temperature levels below 35°C degrees during each treatment. However, this confirmation does not rule out the temperature changes in the microenvironment, due to the inherent abilities of light-absorbing ICG. Prokaryotic cells contain the same heat shock proteins as eukaryotic cells, enabling bacteria to cope against hostile environments. Heat stress has been shown to cause a distinct response in the *S. mutans* expression profile of multiple regulator and other functional genes[22,23]. The extracellular matrix architectural structure and the cells ability to bind are impaired due to heat. Glycosyltransferase (gtf)-c, which is responsible for generating only partially water-soluble glucan, is upregulated; but gtf-b, which is responsible for producing the water-soluble external environment, is not. This change is detectable as early as five to ten minutes after the heat exposure. Similar early responses in the *S. mutans* expression profile can be seen in other heat responsive genes, such as grpE, dnaK and fruR. Eventually, the upregulation of clpE and clpP aid the survival of the cells in the harsh conditions, including increased oxidative stress [23]. Antibacterial blue light has also been shown to alter the gene expression of *S. mutans*, upregulating several genes such as gtfB, brp, smu630, and comDE, but has also been shown to increase the susceptibility of bacteria to ROS [24], which can further explain not only the adaptive mechanisms but also the additive bactericidal effect of the simultaneous use of aBL and aPDT.

We used spectroscopy to measure the ICG absorption properties in the *S. mutans* solution. To our knowledge, no previous work has been published to investigate this. Indocyanine green has been previously shown to undergo redshift of the absorption maximum from 780 nm in water to 805 nm in plasma upon binding to albumin in blood plasma or when binding to lipid structures [25,26]. We found that a similar redshift was also present in planktonic *S. mutans* solution, providing evidence of ICG adhering to bacterial proteins and membrane structures. The lower intensity peak was due to the higher ICG concentration in water compared to that of *S. mutans*-bound ICG. The antibacterial effect of the adhered ICG was estimated by subsequent light excitation, with an 810-nm LED light source resulting in total bacteria killing with light intensities from or above 20J/cm^2^. These results prove that the antibacterial activity is caused by the bacterial-bound ICG and not by the water-solubilized ICG (data not shown). The antibacterial action of ICG was preserved in the bound ICG despite the lower ICG concentration, indicating the role of ICG binding to bacteria as the key treatment-targeting mechanism.

The ability of *S. mutans* to cope with ICG/810-nm aPDT was effectively eliminated by adding simultaneous aBL to the treatment. Antibacterial blue light was markedly less bactericidal than aPDT when similar light doses were compared, but aBL improved the antibacterial effect of aPDT and brought with it the important function of providing a sustained effect in repeated antibacterial treatment. The mechanism is an issue yet to be resolved. In addition to the possible stress response, the ability of the *S. mutans* biofilm external structure reformation to inhibit ICG penetration into the biofilm is surpassed by the ability of aBL to penetrate deep into the layers of the biofilm and inhibit EPS matrix formation, increasing the overall exposure of the bacterial population [12]. Thus, the limitations of the aPDT and photosensitizer penetration into the biofilm could be overcome by dual-light administration. To sum up, results from the present experiments open up new avenues for hypothesis generation and, more practically, for developing devices for biofilm control, especially in preventive dentistry.

